# Broad-Spectrum Coronavirus Inhibitors Discovered by Modeling Viral Fusion Dynamics

**DOI:** 10.1101/2024.03.28.587229

**Authors:** Charles B. Reilly, Joel Moore, Shanda Lightbown, Austin Paul, Sylvie G. Bernier, Kenneth E. Carlson, Donald E. Ingber

**Affiliations:** Wyss Institute for Biologically Inspired Engineering at Harvard University, Boston, MA 02115, USA; Harvard John A. Paulson School of Engineering and Applied Sciences, Harvard University, Cambridge, MA 02138, USA; Vascular Biology Program and Department of Surgery, Harvard Medical School and Boston Children’s Hospital, Boston, MA 02115, USA

**Keywords:** Broad-spectrum, antiviral, molecular-dynamics, coronavirus, AI

## Abstract

Broad-spectrum therapeutics capable of inhibiting SARS-CoV-2, its variants, and related coronaviruses hold promise in curbing the spread of COVID-19 and averting future pandemics. Here, we employed a multidisciplinary approach that included molecular dynamics simulation (MDS) and artificial intelligence (AI)-based docking predictions to identify potent inhibitors that target a conserved region within the SARS-CoV-2 spike protein that mediates membrane fusion by undergoing large-scale mechanical rearrangements. *In silico* binding screens honed in on this region, leading to the discovery of FDA-approved drugs and novel molecules predicted to disrupt spike protein conformational changes. These compounds significantly inhibited SARS-CoV-2 infection and blocked the entry of spike protein-bearing pseudotyped α, β, γ, δ variants as well as SARS-CoV and MERS-CoV in cultured human ACE2-expressing cells. The optimized lead compound significantly inhibited SARS-CoV2 infection in mice when administered orally.

## Introduction

Coronaviruses, including the common cold, cause over 30% of all respiratory tract infections in humans (*1*), including the common cold. They also can lead to widespread illness and mortality worldwide, as we have learned from the emergence of SARS-CoV, MERS-CoV, and now SARS-CoV-2, which has led to the current Coronavirus Disease 2019 (COVID-19) pandemic. While vaccination for SARS-CoV-2 can be protective, acceptance of this therapeutic modality is not widespread. Moreover, throughout the COVID-19 pandemic, SARS-CoV-2

α, β, γ, δ and ο variants of concern (VOC) emerged that can have higher infectivity and increased resistance to vaccines, antibody-based treatments, and targeted antiviral therapeutics (*2*). Therefore, antiviral drugs that exhibit broad-spectrum activity against multiple respiratory coronaviruses are urgently needed, and this will likely be accomplished by identifying new molecular targets (*3–5*). Equally important is the need for oral therapies that can be distributed across populations rapidly and used prophylactically to protect against initial infection, particularly in low-resource nations where vaccines might be costly or challenging to obtain and intravenous administration is difficult.

Attractive drug targets for developing broad-spectrum antiviral therapeutics include viral surface proteins that mediate initial membrane fusion required for entry into host cells, such as the SARS-CoV-2 spike (S) protein. As a result of the COVID-19 crisis, numerous rational drug design and repurposing efforts have targeted this cell surface receptor-binding glycoprotein. Most of these drugs have targeted sites on the external surface of the S protein to interfere with its binding to the host membrane ACE2 receptor (*2,3*), and virtually all therapeutic antibodies target this same interaction site (*6*) or other exposed surface regions on the molecule. However, these same surface binding sites also readily mutate, potentially limiting the spectrum of action and long-term efficacy of available therapeutics (*7–10*). Mutations in different variants were addressed in our study by employing a rational approach using molecular dynamic simulation (MDS) to design drugs aimed at the S proteins’ conserved internal regions, which are crucial for the membrane fusion-dependent viral entry to occur.

Infection begins when the SARS-CoV-2 S protein, a homotrimer (Fig. 1A), binds to ACE2 receptors on the surface of host airway cells. Subsequent cleavage by host proteases (e.g., TMPRSS2, Furin) divides the protein into S1 and S2 (*11–14*). S1 contains the host-receptor binding domain (RBD) and displays vast diversity across different coronaviruses (*15*). In contrast, S2 (and sometimes S’) is mostly buried internally in the pre-fusion state, and it is not subjected to the same level of selective pressure from host immune responses that drive high levels of mutation.

**Figure 1.**
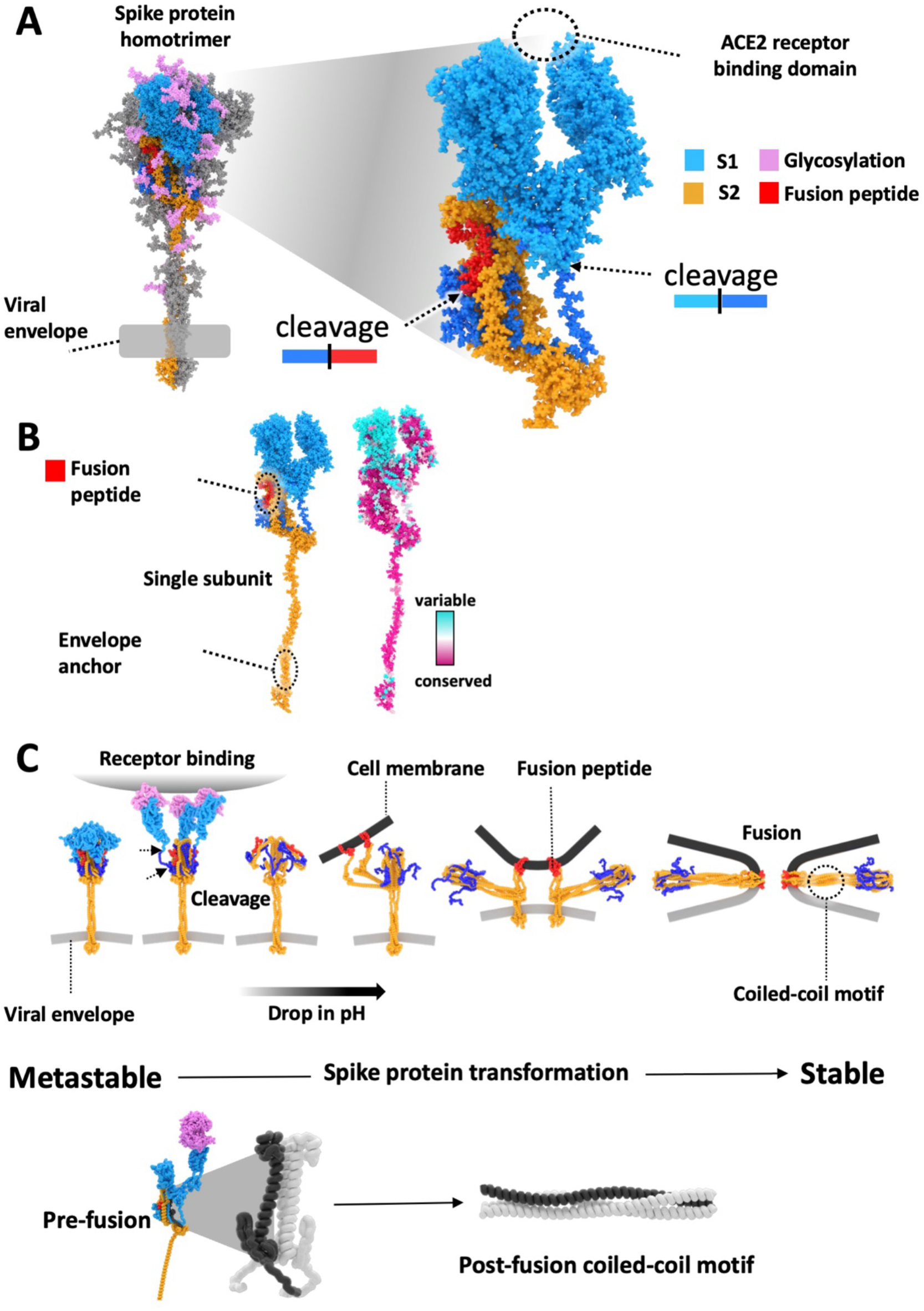
The Spike (S) protein homotrimer facilitates membrane fusion. (**A**) Homology model of the S protein homotrimer in its metastable pre-fusion state. Host protease cleavage sites are indicated along with the fusion peptide (red), S1(blue), and S2 (orange) subunits. (**B**) (left) A single S1 and S2 subunit with the mapping (right) of results from an evolutionary conservation analysis showing evolutionary variation is predominantly found in the vicinity of the receptor-binding domain of the S1 subunit. (**C**) A schematic of the S protein transformation during the viral membrane fusion process as it is thought to proceed (from left to right). The S protein starts in a metastable state and binds to ACE2 proteins on the cell surface. The viral particle is endocytosed while host proteases on the cell surface and the endosome cleave the S protein to generate the fusion peptide and S2 subunit. The S1 units are liberated from the S2 units, and the decreasing pH of the endosome causes the internal rearrangement of S2 structures. As the S2 structures rearrange, the fusion peptides are embedded in the host cell membrane. The S2 structure continues to rearrange to generate a highly stable coiled-coil motif facilitated by heptad repeats found within the S2 structure. The formation of this stable motif drives the trimer into its highly stable post-fusion structural formation, which is irreversible and eventually pulls the viral envelope into contact with the host cell membrane, causing the two lipid bilayers to fuse.

While progress has been made using AI and specialized techniques for potent *in vitro* S protein binders (*16*), effective broad-spectrum therapeutics remain elusive. Our strategy to address this challenge involved an interdisciplinary approach to identify orally bioavailable therapeutics to confront COVID-19 and enable wider therapeutic access amongst the population. We conducted an evolutionary conservation analysis and mapped it to the protein structure (Fig. 1B) to identify a drug target region. This analysis revealed the variability within the S1 protein, which is also reflected in the locations of mutations within VOCs designated by the World Health Organization (WHO) that have emerged throughout the COVID-19 pandemic (Fig. 1B and Fig. S1) (*17*). In contrast, the S2 subunit is less prone to mutation, and it also contains both the internal viral envelope anchoring region and the extracellular fusion peptide sequence that mediates fusion with the host cell membrane.

Past work has shown that the initial binding and cleavage events when the SARS-CoV-2 virus binds to the host cell surface trigger the S protein’s metastable pre-fusion structure to undergo a large-scale mechanical transition (Fig. 1C) (*18*). Fusion of the viral envelope and host membrane, and hence viral entry, is made possible because the S2 subunit forms a highly stable structural conformation resulting from this mechanical transformation that facilitates embedding the fusion peptide regions of S2 within the host cell membrane. Additional microenvironmental triggers, including a drop in pH within membrane vesicles during endocytosis of the bound viral particles, also help drive and stabilize this mechanical state transition (Fig. 1C) (*11*, *19*, *20*). These features, which have been highly conserved throughout evolution, make the S2 subunit an attractive S protein component to use as a drug target for preventing coronavirus entry.

In viral infection, the S2 subunit transitions from a compact pre-fusion to an elongated post-fusion state, stabilized by coiled-coil motifs from heptad repeat (HR) domains (Fig. 1C) (*19*, *21–23*). The structures of these states are known from cryoelectron microscopy (cryoEM) and crystallography, but the transition dynamics remain unclear. Prior therapeutics aimed to halt fusion by targeting static pre-fusion and post-fusion S2 conformations (*24*) or indirectly via protease inhibitors that block the S2 cleavage needed to generate fusion peptides (*25–27*). Our study aims to develop drugs against the transition dynamics using MDS to identify a compelling molecular binding site, potentially hindering S2’s conformational changes required for membrane fusion.

## Results and Discussion

The primary goal of this study was to develop broad-spectrum coronavirus therapeutics by identifying a druggable binding pocket within the S2 subunit that remains hidden in the prefusion state and avoids selective pressure by antibodies and other environmental stimuli that drive high levels of mutation on the surface of proteins. To achieve this, we initially utilized molecular dynamic simulations (MDS) and homology models (*28*) based on SARS-CoV2 sequencing data. This project was initiated early in the COVID-19 pandemic, and as the pandemic progressed, homology models were refined for VOCs (*29*). These simulations incorporated parameters like viral envelope anchorage, receptor binding domain (RBD) pulling due to ACE2 binding, and microenvironmental conditions mimicking host cell interactions which aid viral entry (e.g., variations in protonation state to mimic the low pH of the endosome, proteolytic cleavage of S protein to generate S2). Simulations were analyzed and visually compared to existing crystal structures of S proteins, focusing on the heptad repeat (HR) regions that form the coiled-coil structures during post-fusion states, and we mapped amino acid conservation scores onto the MDS trajectories (Fig. 1B).

This MDS analysis revealed a potential drug-targeting site within the highly conserved secondary structure of the S2 domain. This site on the S2 unit is located where it becomes separated from residues 716-724 (peptide sequence TNFTISVTT) of the S1 domain during host-cell interaction (Fig. 2, A and B). During the simulations, this section of the S1 unit separates when a pulling force is applied to the RBD domain to emulate the tensional forces experienced in response to it binding to the ACE2 receptor on the host cell surface. As the simulation progresses, a cavity forms, exposing highly conserved residues (Fig. 2, A and B) before entering the endosome where the pH is neutral. Mechanical force application to the RBD also caused the release of the fusion peptide from its buried position. Interestingly, these simulations suggested that mechanically induced unpacking of the fusion peptide after binding the RBD to its receptor might be necessary for facilitating the access of proteases to the cleavage site that liberates the fusion peptide.

**Figure 2.**
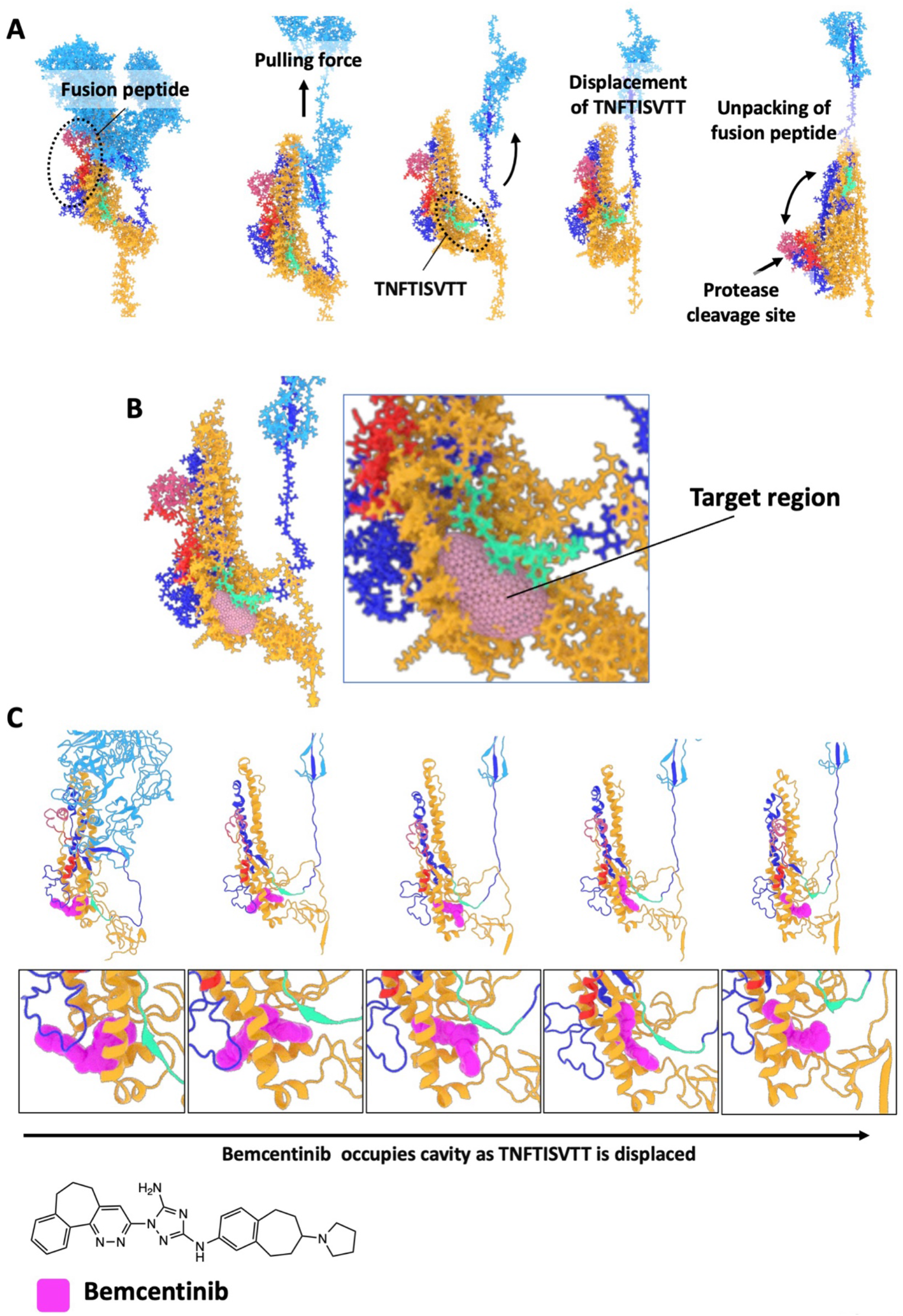
Targeting Spike protein fusion dynamics for Broad Spectrum efficacy against SARS-CoV2. (**A**) Conformations of the S protein subunit taken from a simulation of the homotrimer with pulling forces applied to the ACE2 binding domain. Showing the displacement of the TNFTISVTT peptide from within a conserved region of the S2 domain and the unpacking of the fusion peptide. (**B**) The region of the S2 subunit that is targeted during *in silico* docking screens. **(C)** Poses of bemcentinib were generated using AI-based blind docking using DiffDock across conformations of the S protein during the same pulling simulation in (**A**). The transition of TNFTISVTT out of the target region due to pulling forces is shown from left to right. As TNFTISVTT transitions out, bemcentinib transitions in to occupy the target cavity. The structure of bemcentinib is shown at the bottom.

Notably, deformations and the separation of the TNFTISVTT peptide in response to mechanical stretching upon RBD binding, including structural variations in the HR1 region, persist even after cleavage and removal of the S1 domain and hence, after the applied tensional forces dissipate (Fig. S2A). Importantly, these deformed S2 units also maintain anchorage to the viral envelope. Additional simulations of the cleaved S protein were conducted with protonated histidines to mimic the acidic environment of the late endosome (pH 4.5) that facilitates viral membrane fusion, resulting in the target site becoming more "open." Based on this observation, the hypothesis was formed that identifying small molecules with a high affinity for this highly conserved site (Fig. 2B) could physically stabilize the region occupied by TNFTISVTT in the pre-fusion state of the intact Spike protein. The concept was that this binding interaction could potentially prevent the deformations caused by the pulling and removal of TNFTISVTT and prevent the large-scale shape shifts required for the S2 subunit to transition to the irreversible post-fusion state, thereby inhibiting membrane fusion and viral entry.

Initial *in silico* docking screens using AutoDock Vina (*30*) were performed against the target pocket to identify potential candidate small molecule inhibitors. Docking receptors were defined based on conformations taken from the MDS trajectory, explicitly focusing on the most significant deformations observed during the first 100 ns of simulations where the protonation states were adjusted to mimic the acidified environment, and the simulations were conducted with the S1 section still occupying the active site and no mechanical forces applied. Subsequently, the S1 section was removed post-simulation to generate conformations where the docked small molecules could mimic the role of the S1 section that typically fills the target pocket in the prefusion state.

This study was initiated soon after the advent of COVID-19 with emergency support from the Defense Advanced Research Projects Agency (DARPA) to leverage our computational MDS approach to rapidly repurpose drugs for the emerging pandemic. Given the urgency, initial screens were performed using the DrugBank library of approved and investigational compounds (*31*) in the hope that we could identify a drug repurposing opportunity. The ∼10,000 compounds within the Drugbank library were then ranked based on the strongest average binding affinity for the top five poses across ten conformations of the target pocket selected every ten ns from the first 100 ns of MDS. The highest-ranked molecule that was also orally bioavailable was the investigational AXL kinase inhibitor drug, bemcentinib. The MDS and docking studies revealed that bemcentinib binds with high affinity to the target region across multiple protein conformations of S2.

Advanced AI techniques emerged throughout the pandemic and were integrated into our *in silico* pipeline. We performed AI-based blind docking predictions, where a target region is not explicitly selected, and used DiffDock (*32*) to assess the potential preferential binding of bemcentinib to our target site throughout the simulation trajectory with pulling forces. We found that as pulling forces initiate displacement of S1-TNFTISVTT from the target pocket, bemcentinib exhibits progressively improved orientations, optimal occupancy, and effective binding within the target site (Fig. 2C).

This observation suggests that bemcentinib’s affinity increases progressively during the simulation in response to the applied pulling force, which should disrupt initial conformational changes triggered by mechanical forces generated by RBD binding to cell surface receptors before proteolytic cleavage and the decrease in pH. Such inhibitory capability is particularly advantageous in light of recent findings that illustrate multiple cellular entry routes for the virus (32). Some of these routes involve endocytosis, while others are independent of it, not requiring the acidic conditions of the late endosome to facilitate viral fusion.

The predicted antiviral activity of bemcentinib against SARS-CoV-2 was confirmed by showing that bemcentinib potently inhibited SARS-CoV-2 infection of human lung A549 cells stably expressing human ACE2 (A549-hACE2) with an IC_50_ of 0.070 μM (Fig. 3A and Table S1). This result is consistent with a recently published study showing that bemcentinib inhibits SARS-CoV-2 infection of human lung Calu-3 cells (*33*). In addition, we confirmed that bemcentinib exerted this antiviral activity by preventing viral entry as it potently inhibited the ability of a VSV-based pseudovirus expressing the SARS-CoV-2 S protein (SARS-CoV-2pp) to infect HEK-293 cells stably expressing human ACE2 (HEK-293-hACE2) with an IC_50_ of 5.7 μM (Fig. 3A and Table 1). As previously demonstrated, we also found that bemcentinib potently inhibits purified AXL kinase with an IC_50_ of 0.9 nM (Fig. S3 and table S1).

**Figure 3.**
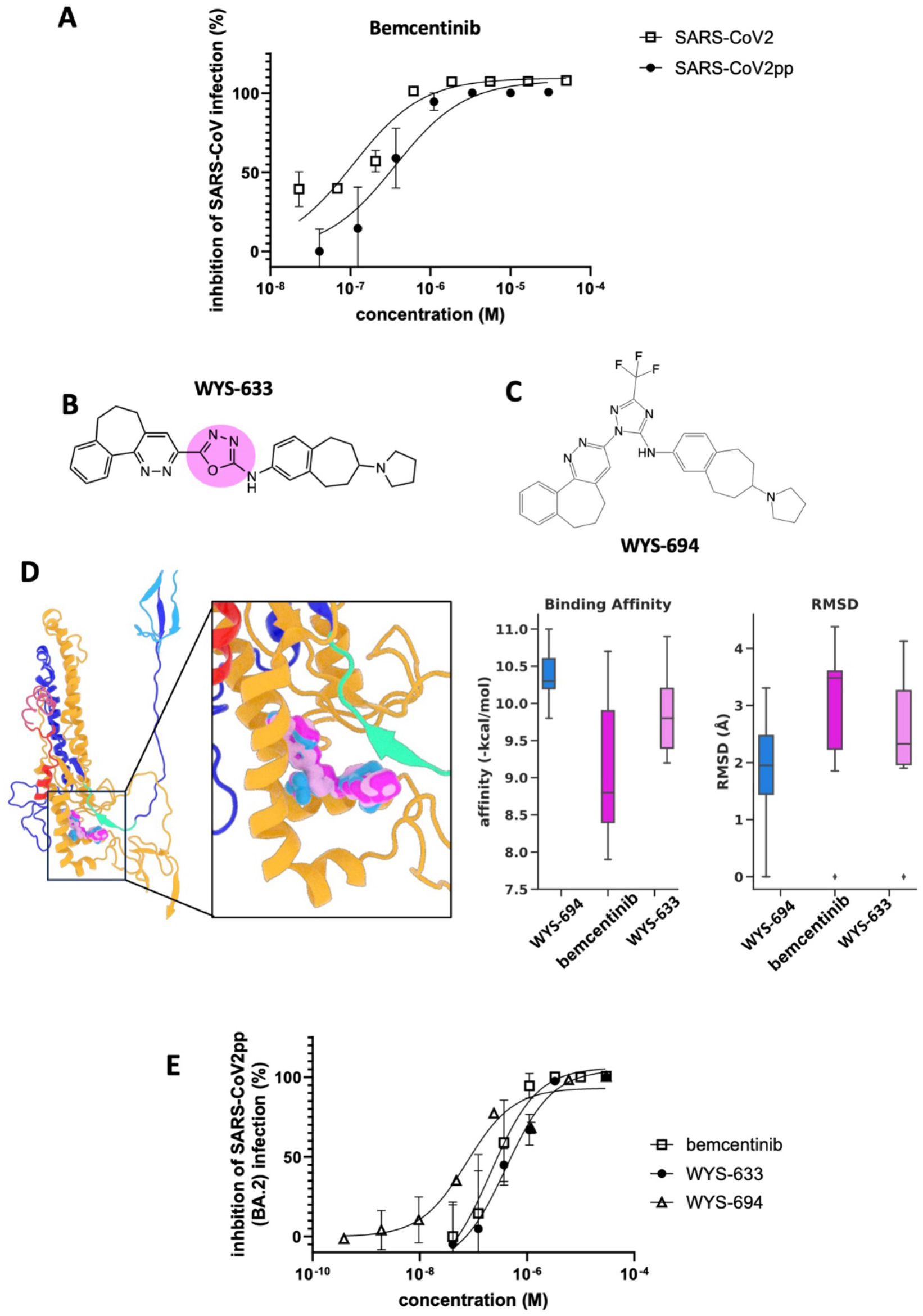
Bemcentinib and novel analogs inhibit infection *in vitro*. **(A)** Effects of bemcentinib against SARS-CoV-2_GFP infected ACE2-expressing A549 cells and BA.2 omicron SARS-CoV-2pp infected ACE2-expressing HEK293 cells. (B)The structure of WYS-633 with the region modified in bemcentinib is highlighted. (C) The structure of WYS-694. (D) Bemcentinib, WYS-633, and WYS-694 docked in the target region. The binding affinity and RMSD analysis of the top 9 poses docked with the S protein conformation shown and scored with smina. (E) Effects of compounds on pseudotyped SARS-CoV-2 viral entry in ACE2-expressing HEK293 cells. The plot shows the inhibitory effects of bemcentinib, WYS-633, and WYS-694 on ACE2-expressing HEK293 cells infected with SARS-CoV-2pp for 48 h. The number of pseudoparticles in the infected cells was quantified by measuring luciferase activity; viral entry in untreated cells was set as 100%.

**Table 1.** Compound EC_50_ values on pseudotyped SARS-CoV-1, SARS-CoV-2 viral infection in ACE2-expressing HEK293 cells or on pseudotyped MERS viral entry in DPP4-expressing HELA cells.

|  | bemcentinib |  | WYS-633 |  | WYS-694 |  |
| --- | --- | --- | --- | --- | --- | --- |
|  | EC<sub>50</sub> (μM) | n | EC<sub>50</sub> (μM) | n | EC<sub>50</sub> (μM) | n |
| SARS-CoV-1pp | 0.91 ± 0.01 | 2 | 0.75 ± 0.07 | 2 | 0.82 ± 1.02 | 3 |
| SARS-CoV-2pp | 5.68 ± 3.76 | 5 | 2.52 ± 2.26 | 7 | 0.08 ± 0.03 | 3 |
| B.1.1.7 (alpha) SARS-CoV-2pp | 0.43 ± 0.18 | 3 | 1.23 ± 0.58 | 3 | 0.12 ± 0.07 | 3 |
| B.1.351 (beta) SARS-CoV-2pp | 0.39 ± 0.14 | 3 | 1.25 ± 0.05 | 3 | 0.18 ± 0.11 | 3 |
| B.1.617.2 (delta) SARS-CoV-2pp | 0.38 ± 0.14 | 3 | 1.05 ± 0.37 | 3 | 0.13 ± 0.09 | 3 |
| B.1.1.28 (gamma) SARS-CoV-2pp | 0.42 ± 0.12 | 3 | 1.66 ± 0.71 | 3 | 0.16 ± 0.15 | 3 |
| B.1.1.529 (omicron) SARS-CoV-2pp | 0.21 ± 0.01 | 2 | 1.03 ± 0.65 | 3 | 0.13 ± 0.12 | 3 |
| BA.2 SARS-CoV-2pp | 0.16 ± 0.03 | 3 | 0.68 ± 0.29 | 3 | 0.45 ± 0.44 | 3 |
| MERSpp | 0.09 ± 0.01 | 2 | 0.31 ± 0.12 | 2 | ND |  |

Values are means ± SD of data from 2 to 7 independent experiments (ND: not determined).

Independent of our ongoing work, bemcentinib was selected for testing in a COVID-19 clinical trial (*34*) based on its *in vitro* antiviral activity against SARS-CoV-2 and the hypothesis that its antiviral activity is dependent on its ability to inhibit AXL kinase (*35*). The antiviral activity of bemcentinib against SARS-CoV-2 was borne out by recent results from a Phase 2 clinical trial in which bemcentinib improved clinical response and key secondary endpoints in patients hospitalized with COVID-19 (*36*).

To determine whether bemcentinib’s ability to inhibit SARS-CoV-2 infection could be the result of the S2 protein interaction predicted by our MDS, we designed and synthesized structurally-similar analogs that maintained preferential *in silico* binding to the S protein target region but lacked key molecular features that would be critical for AXL kinase inhibition. The lead molecule from this chemical series, WYS-633, dose-dependently inhibited SARS-CoV-2 infection of A549-hACE2 cells with an IC_50_ of 0.61 μM (Table S1) and appeared to act at the level of viral entry by potently inhibiting SAR-CoV-2pp infection of HEK-293-hACE2 with an IC_50_ of 2.52 μM (Fig. 3E and Table 1). Notably, the ability of WYS-633 to inhibit SARS-CoV-2 infection was independent of AXL kinase inhibition as the compound was completely inactive against AXL kinase, as we had predicted (Fig. S3 and Table S1). Bemcentinib is also known to disrupt lysosomal function and cause significant vacuolization in cells, independent of its AXL kinase activity (*37*). This lysosomal disruption is another possible mechanism by which this molecule could produce antiviral activity. However, while bemcentinib induced significant vacuolization in human lung A549 cells, WYS-633 did not (Fig. S4). To further clarify the mechanism of action of WYS-633, we also investigated the potential role of phospholipidosis as an agency of antiviral activity. It has recently been shown that phospholipidosis, specifically as it relates to cationic ampiphilic drugs (CADs), is a shared mechanism of antiviral activity within drug repurposing screens (*38*). However, while both bemcentinib and WYS-633 are cationic ampiphilic molecules, neither compound demonstrated phospholipidosis when tested for this effect in A549 cells using the well-established NBD-PE staining assay (*38*) in which amiodarone was used as the positive control (Fig. S5). These results obtained with the bemcentinib analog, WYS-633, demonstrate that this molecule inhibits SARS-CoV2 entry into cells in an AXL kinase- and vacuolization-independent manner. They are also consistent with our hypothesis that this molecule directly binds the S protein, as our computational models predicted.

It is important to note that the conserved target pocket we identified in the SARS-CoV-2 spike protein is also found within related S proteins on the surfaces of all SARS-CoV-2 VOCs identified as well as on other coronaviruses, such as SARS-CoV and MERS. Therefore, we tested our novel analog in infection assays using a panel of pseudotyped viruses expressing these different S protein variants. WYS-633 inhibited entry of SARS-CoV-1, MERS, and the SARS-CoV-2 VOCs (α, β, δ, γ, ο and the BA.2 ο variant) with approximately equal high nM to low μM potency (Table 1), demonstrating the broad-spectrum activity of this compound.

New analogs of WYS-633 were designed and synthesized to improve antiviral efficacy and drug-like properties. A standout among these was WYS-694 (Fig. 3C), which has altered molecular connectivity in which the key flanking groups adopt a 1,2-positioning instead of the bemcentinib-like 1,3-positioning. This molecule exhibited docking preference for the target region (Fig. S2B) with significantly higher *in silico* binding affinity and lower root mean square deviation (RMSD) compared to bemcentinib and WYS-633 when the binding affinities of the top 9 poses for each docked molecule were evaluated using smina docking and scoring (*39*) (Fig. 3D). As the RMSD quantifies how much the lower-ranked poses deviate from the highest-ranked pose, these results indicate that WYS-694 exhibits a significantly stronger binding affinity for the target region than either bemcentinib or WYS-633. WYS-694 also proved to be on average 12.5 fold more potent than WYS-633 in inhibiting viral entry of wild-type SARS-CoV-2 pseudoviruses (Fig. 3E) and a panel of pseudotyped viruses expressing SARS-CoV-2 VOCs (Table 1).

While designing new analogs, we aimed to improve drug-like properties, including optimizing pharmacokinetic properties. This is reflected in pharmacokinetic experiments conducted in mice (Table 2). Bemcentinib (100 mg/kg) demonstrated significant oral bioavailability with a high maximal plasma concentration (C_max_ = 1,757 ng/ml) and a prolonged time to reach maximal drug concentration (T_max_ = 13 hours), resulting in substantial plasma exposure over time or area under the curve (AUC = 33,287 h*ng/ml). WYS-633 at the same dose, showed oral bioavailability with a reasonable C_max_ but had a shorter T_max_ (2 hours), leading to lower overall exposure. In contrast, our optimized analog, WYS-694 (30 mg/kg) exhibited similar initial exposure to WYS-633 but an extended T_max_ (24 hours), resulting in significantly improved drug exposure over time.

**Table 2.** Pharmacokinetics of bemcentinib and novel analogs in mice. Bemcentinib and WYS-633 were formulated in 0.5% hydroxypropyl methylcellulose and 0.1% Tween 80 and WYS-694 was formulated in 100% PEG300 and administered at the indicated doses by oral gavage in C57 Bl/6 mice. Blood was drawn at 0.25, 0.5, 1, 2, 4, 8 and 24 hours and drug concentrations in plasma were determined by LC-MS/MS. Exposure levels over time were graphed and from the exposure profile Tmax, CMax and AUC (area under the curve) were determined.

| Test Article | Dose<br>(mg/kg) | TMax<br>(h) | CMax<br>(ng/ml) | AUC<br>(h*ng/ml) |
| --- | --- | --- | --- | --- |
| Bemcentinib | 100 | 13 | 1757 | 33287 |
| WYS-633 | 100 | 2 | 950 | 7921 |
| WYS-694 | 30 | 24 | 556 | 10698 |

With the pharmacokinetic profiles of these molecules determined, their *in vivo* antiviral activity was assessed in SARS-CoV2-infected mice that overexpress human ACE2 (Fig. 4). Bemcentinib, WYS-633, and WYS-694 were dosed prophylactically at 24 hours and then 2 hours before intranasal administration of SARS-CoV2 virus followed by a final dose of compound at 24 hours post-infection. Viral load was determined in lung homogenates three days post-infection. When dosed at 100 mg/kg p.o., bemcentinib failed to reduce SARS-CoV2 viral load despite having a significant and prolonged exposure (Fig. 4B). WYS-633 dosed at 100 mg/kg p.o., similarly failed to inhibit SARS-CoV2 infection (Fig. 4C). In contrast, when WYS-694 was dosed at 30 mg/kg p.o. to match the overall exposure of WYS-633, it significantly inhibited SARS-CoV-2 infection and reduced viral load in these mice by more than 4-fold (Fig. 4D).

**Figure 4.**
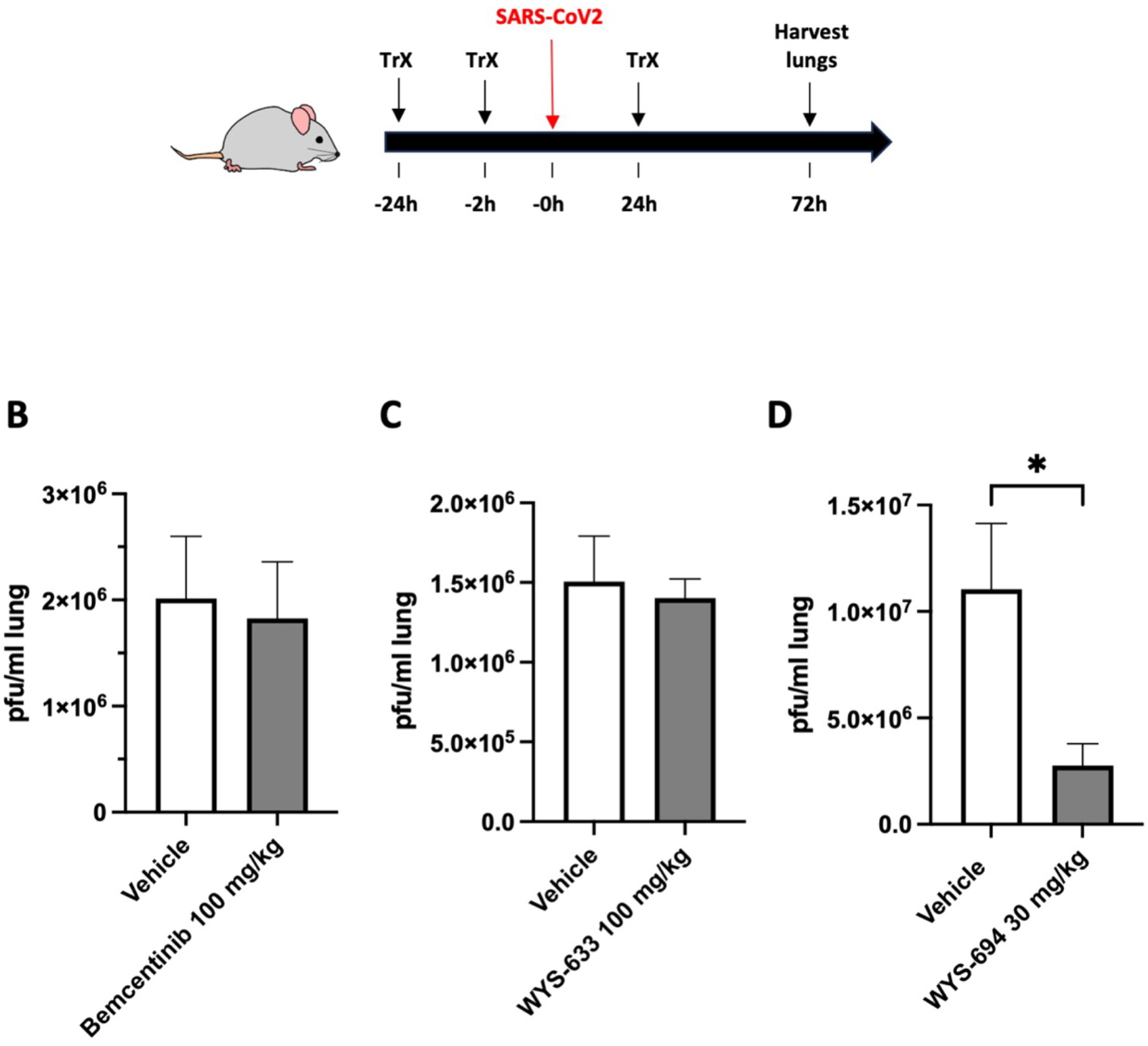
Efficacy of bemcentinib and novel analogs to inhibit SARS-CoV2-infection in mice that overexpress human ACE2. (A) Antiviral efficacy of small molecule compounds *in vivo* was assessed in K18 ACE2 over-expressing mice by oral gavage (“Trx”) of compounds at 24 hours before, 2 hours before, and 24 hours after intranasal administration of 2 x 10^4^ pfu/mouse of SARS CoV-2 WA1/2020. After three days, viral load was determined in lung homogenates by plaque assay in Vero-6 cells and was expressed as pfu/ml lung homogenate for (B) Bemcentinib was formulated in 0.5% hydroxypropyl methylcellulose and 0.1% Tween-80 and dosed at 100 mg/kg p.o. (C) WYS-633 was formulated in 0.5% hydroxypropyl methylcellulose and 0.1% Tween-80 and dosed at 100 mg/kg p.o., and (D) WYS-694 was formulated in 100% PEG300 and dosed at 30 mg/kg p.o. to match the overall exposure of WYS-633.

Traditional approaches to combatting COVID-19 have typically focused on developing new drugs targeting the S protein’s surface for ACE2 binding, and contemporary AI techniques seek to replicate traditional drug-targeting approaches. In contrast, with DARPA’s support to find repurposable drugs, we adopted a novel strategy utilizing molecular dynamic simulations (MDS) to analyze the dynamic mechanical changes in the SARS-CoV-2 S protein necessary for membrane fusion and viral entry. We developed a drug pipeline to rapidly integrate nascent computational approaches and combine these with traditional medicinal chemistry strategies. This approach uncovered bemcentinib, an orally bioavailable drug, which appears to directly bind a conserved region within the S2, undergoing critical shape changes for membrane fusion. While previous studies suggested bemcentinib’s efficacy against SARS-CoV-2 was due to AXL kinase inhibition and/or lysosomal interference, our novel analog synthesis led to a more potent compound that lacks these activities yet still effectively inhibits SARS-CoV-2 infection *in vitro* and *in vivo*.

Moreover, because the target pocket we identified in the S2 protein is highly conserved, these new compounds exhibit broad-spectrum antiviral activity and inhibit S protein-dependent entry of 5 different SARS-CoV-2 VOCs as well as SARS-CoV and MERS-CoV. Thus, this novel compound represents the first in a new class of antiviral compounds that target mechanical transformations within viral S proteins. This compound offers an alternative approach to potentially prevent SARS-CoV-2 infection and treat COVID-19 patients, and provides a new tool to counter novel coronaviruses that will likely emerge in the future. This interdisciplinary approach, combining computational MDS with AI and medicinal chemistry, also presents an exciting opportunity to discover drugs that target other virus classes that utilize membrane fusion proteins (e.g., influenza, HIV, Ebola, Marburg, Nippah, Dengue, Measles, and more).

## Materials and Methods

### Molecular dynamics simulation

Initial homology models of the spike protein were generated utilizing pdb: 6crz,6xra,6vsb and 6vxx with the assistance of the Modeler software and using the sequence with uniprot id: P0DTC2. Glycosylation was incorporated using the Charm GUI(40). Protonation states were determined through the H++ server(41) at pH levels of 4.5 and 7. Molecular dynamics simulations, employing both implicit and explicit solvents, were performed with the Amber forcefield using Ambertools(42). These simulations were executed utilizing OpenMM(43). Subsequent analysis of the simulation data was conducted using the MDtraj toolkit(44).

### Evolutionary conservation analysis

To evaluate the evolutionary conservation of specific residues within the spike protein, we performed a conservation analysis using the ConSurf webserver(15)l. Subsequently, we utilized SideFX Houdini to map these conservation scores onto the molecular structure, enabling a visual representation of the conserved regions within the protein.

### *In Silico* Docking

For high-throughput in silico docking screens, we employed Vina(30), utilizing conformations generated with MDtraj extracted from MD trajectories. Additionally, we utilized Diffdock(32) for blind artificial intelligence-based docking. Furthermore, Smina(39) was employed for docking studies based on the binding regions identified through the Diffdock analysis.

### Visualizations

Procedural modelling and visualization were carried out using VMD, MDTraj, python and SideFX Houdini.

### Infection assay using pseudotyped viruses in ACE2-expressing HEK293 cells

Compounds were tested using entry assays for SARS-CoV-2 pseudoparticles (SARS-CoV-2pp), as previously described (Longlong et al paper). SARS-CoV-2pp and its variants B.1.1.7 (alpha), B.1.351 (beta), B.1.617.2 (delta) and B.1.1.28 (gamma)) were obtained from Cellecta Inc and B.1.529 (omicron) and BA.2 were obtained from Virongy. Infections were performed in 96-well plates. Pseudotyped viruses were added to 20 000 ACE2-expressing HEK293 cells per well in the presence or absence of the test compound. The mixtures were then incubated for 48 hours at 37°C. Luciferase activity, which reflects the number of pseudoparticles in the host cells, was measured at 48 h post-infection using the Bright-Glo reagent (Promega) according to the manufacturer’s instructions. Test drug was serially diluted to a final concentration of 0-30 μM. The maximum infectivity (100%) was derived from the untreated wells; background (0%) from uninfected wells. To calculate the infection values, the luciferase background signals were subtracted from the intensities measured in each of the wells exposed to drug, and this value was divided by the average signals measured in untreated control wells and multiplied by 100%.

### Infection assay using MERS pseudotyped viruses in DPP4-expressing HELA cells

Compounds were tested using entry assays for MERS pseudoparticles (MERSpp). MERSpp was obtained from Cellecta Inc. Infections were performed in 96-well plates. Pseudotyped viruses were added to 7 500 DPP4-expressing HELA cells per well in the presence or absence of the test compound. The mixtures were then incubated for 48 hours at 37°C and luciferase activity assay was performed as described above.

### A549 cell vacuolization

A549 cells were obtained from American Type Culture Collection (catalog # CCL-185) and cultured in DMEM medium supplemented with 10% of FBS. Cells were plated in culture medium at 150 000 cells per well in 24-well plates one day before use. Cells were incubated with DMSO (vehicle) or 5 µM of compounds for 24 hours. Phase-contrast images were captured with Echo Revolve.

### A549 cell phospholipidosis

A549 cells were plated in culture medium at 15 000 cells per well in 96-well plates one day before use. The assay was performed as previously described (*10*). Briefly, cells were treated for 24 hours with a dose-range of different compounds in presence of 7.5 μM NBD-PE (1-Palmitoyl-2-[12-[(7-nitro-2-1,3-benzoxadiazol-4-yl)amino]dodecanoyl]-sn-Glycero-3-phosphoethanolamine, (ThermoFisher, cat # N360)). Amiodarone (Sigma, cat# A8433) was used as a positive control for phospholipidosis. The plated cells were washed twice with Hanks’ Balanced Salt Solution (HBSS, Gibco cat # 14025-092), and 50 μL of HBSS was added to each well. Fluorescence of NBD-PE taken up by cells was measured with a fluoromicroplate reader (Biotek, Synergy H1) with an excitation wavelength of 485 nm and emission of 538 nm. After measuring the fluorescence of NBD-PE, 50 μL of HBSS containing 20 μg/mL of Hoechst33342 was added to measure total cell population. Incubation was continued for 20 minutes at 37°C, after which the fluorescence was measured with an excitation wavelength of 355 nm and emission of 460 nm. Normalized values were calculated by dividing the NBD-PE value by the Hoechst33342 value.

### AXL kinase assay

AXL kinase enzyme potencies were determined by Reaction Biology (www.reactionbiology.com) using their Hot Spot kinase Assay. The base reaction buffer for the assay was 20 mM Hepes (pH 7.5), 10 mM MgCl_2_, 1 mM EGTA, 0.01% Brij35, 0.02 mg/mL BSA, 0.1 mM Na_3_VO_4_, and 2 mM DTT with a 1% DMSO concentration. Required cofactors were added individually to each kinase reaction. The substrate was freshly prepared in the reaction buffer described above, and then cofactors were delivered. The purified kinase was added to the substrate solution and then gently mixed. Compounds were added from 100% DMSO into the kinase reaction mixture by Acoustic technology (Echo550; nanoliter range) and then incubated for 20 min at room temperature. ^33^P-ATP (10 µM) was delivered to initiate the reaction, and then the mixture was incubated again for two hours at room temperature. Kinase activity was determined by P81 filter-binding method as described in the following reference: Anastassiadis T, *et al*. Comprehensive assay of kinase catalytic activity reveals features of kinase inhibitor selectivity. *Nat. Biotechnol*. 2011 Oct 30;29(11):1039-45. doi: 10.1038/nbt.2017

### Pharmacokinetics in mice

Bemcentinib and WYS-633 were formulated in 0.5% hydroxypropyl methylcellulose and 0.1% Tween 80 and WYS-694 was formulated in 100% PEG300 and administered to C57 Bl/6 mice (n=3 per group) at 100 mg/kg (Bemcentinib, WYS-633) or 30 mg/kg (WYS-694) by oral gavage. Blood samples were drawn at 0.25, 0.5, 1, 2, 4, 8 and 24 hours and plasma was prepared. The desired serial concentrations of working solutions were achieved by diluting stock solution of analyte with 50% acetonitrile in water solution. 5 µL of working solutions (1, 2, 4, 10, 20, 100, 200, 1000 and 2000 ng/mL) were added to 10 μL of the blank C57BL/6J mice plasma to achieve calibration standards of 0.5∼1000 ng/mL (0.5, 1, 2, 5, 10, 50, 100, 500 and 1000 ng/mL) in a total volume of 15 μL. Five quality control samples at 1 ng/mL, 2 ng/mL, 5 ng/mL, 50 ng/mL and 800 ng/mL for plasma were prepared independently of those used for the calibration curves. These QC samples were prepared on the day of analysis in the same way as calibration standards.15 μL standards, 15 μL QC samples and 15 μL unknown samples (10 µL plasma with 5 µL blank solution) were added to 200 μL of acetonitrile containing IS mixture for precipitating protein respectively. Then the samples were vortexed for 30 s. After centrifugation at 4 degree Celsius, 3900 rpm for 15 min, the supernatant was diluted 3 times with water. 10 µL of diluted supernatant was injected into the LC/MS/MS (AB API 5500+ LC–MS/MS instrument) with a HALO C18 90A 2.7µm (50*2.1 mm) column) for quantitative analysis. The mobile phases used were 95% water (0.1% formic acid) and 95% acetonitrile (0.1% formic acid). Data was graphed as exposure levels over time from which TMax, CMax, and AUC (area under the curve) were determined. All PK studies were conducted by Pharmaron and performed in accordance with the guidelines of the Institutional Animal Care and Use Committee of Pharmaron.

### Efficacy studies in SARS-CoV2 infected mice

Bemcentinib and WYS-633 were formulated in 0.5% hydroxypropyl methylcellulose and 0.1% Tween 80 and WYS-694 was formulated in 100% PEG300 and administered to K18-hACE2 female mice (Jackson Labs) at 100 mg/kg (Bemcentinib, WYS-633) or 30 mg/kg (WYS-694) by oral gavage (n=6 per group). Compounds were administered at 24 hours before, 2 hours before, and 24 hours after intranasal administration of 2 x 10^4^ pfu/mouse of SARS CoV-2 2019-nCoV/USA-WA1/2020 (1/2 dose per nare). Mice were weighed daily, body temperatures were measured, and mice were physically assessed for clinical signs. All animals were sacrificed at 3 days post infection, and the lungs were collected for analysis. The right lobe was collected for viral titer determination and left lobe was collected in formalin for fixation and inactivation. Viral titer was determined by plaque assay. Twelve-well plates were seeded with Vero E6 cells overnight. A series of 10-fold dilutions of lung homogenate were made in MEM with 2% FBS, and 200 µL of each dilution was added to each well in duplicate and incubated at 37°C, 5% CO2 for 1 h. Following incubation, overlay media consisting of a 1:1 mix of 2% carboxymethylcellulose and 2XMEM supplemented with 10% FBS and 2% penicillin-streptomycin was added to each well. Plates were incubated at 37°C, 5% CO2 for 72 h. Wells were fixed with 10% formalin, stained with 1% crystal violet, washed, and plaques were counted. All efficacy studies were conducted at the Regional Biocontainment Laboratory at the University of Tennessee Health Science Center and performed in accordance with the guidelines of their Institutional Animal Care and Use Committee.

## Funding

Defense Advanced Research Projects Agency (HR0011-20-2-0040), Open Philanthropy – Good Ventures Foundation, and Wyss Institute for Biologically Inspired Engineering at Harvard University.

## Notes

### Competing Interest Statement

The authors have declared no competing interest.

