## Supplementary material for "Broad-Spectrum Coronavirus Inhibitors Discovered by Modeling Viral Fusion Dynamics": SI appendix

**This PDF file includes:**

Figures S1 to S5
Tables S1 to S2

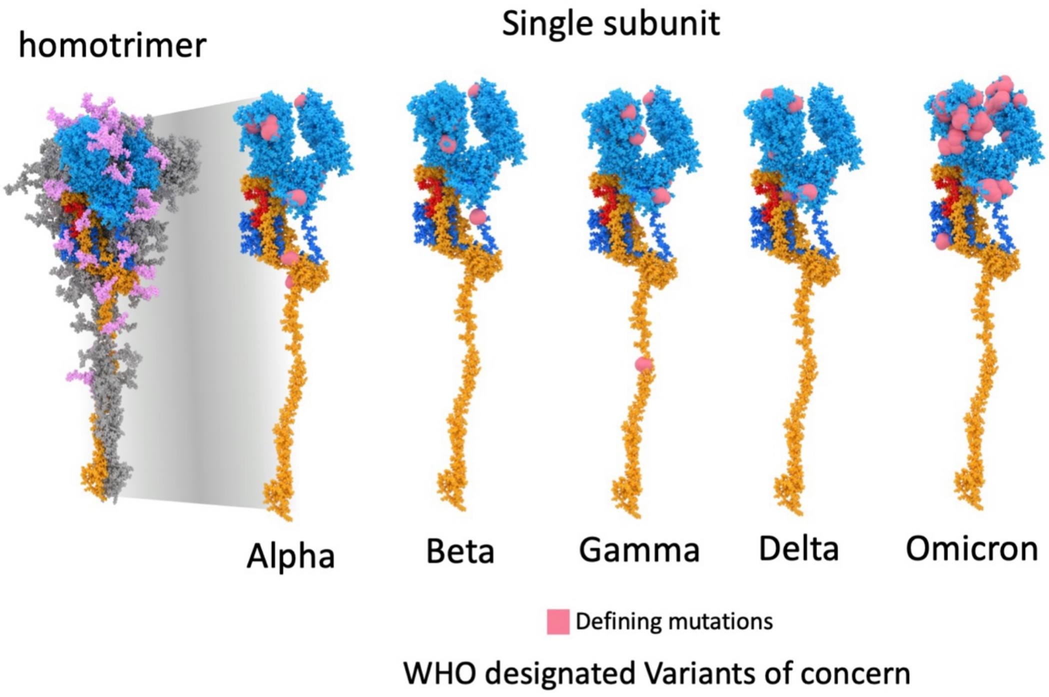


**Figure S1. Location of defining mutations within the S protein for WHO designated variants of concern**. Alpha ( B.1.1.7), Beta ( B.1.351), Gamma (P.1), Delta (B.1.617.2), Omicron (B.1.1.529)**.**

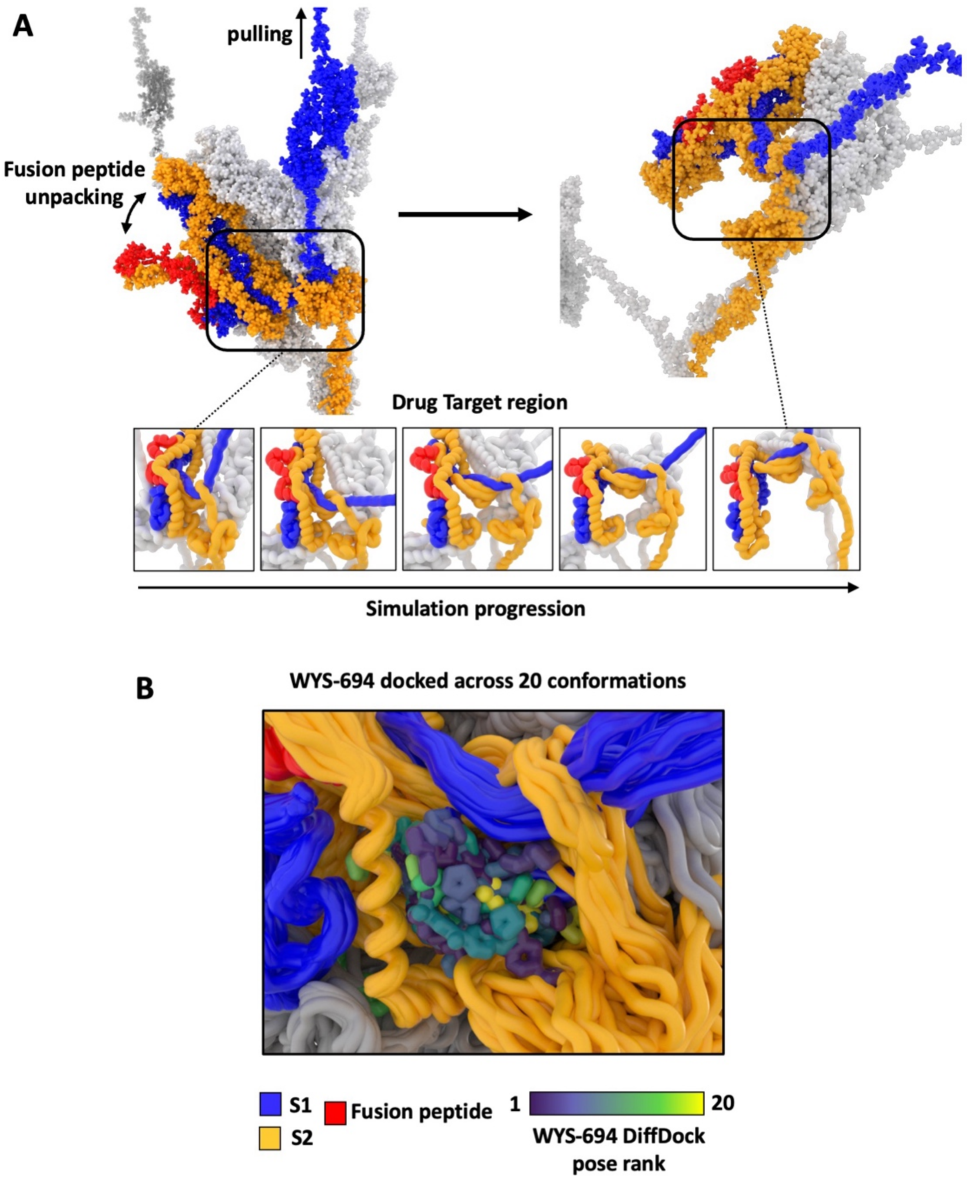


**Figure S2. Drug target region during molecular dynamic simulation.** (A) Simulation of spike protein under acidic conditions, pulling forces and envelope anchorage. (B) 20 conformations were selected from across the simulation in (A), DiffDock poses are shown for WYS-694 for each conformation and rendered based on rank for each target conformation. Target and pose are aligned to the heptad repeat domain and superimposed for rendering.

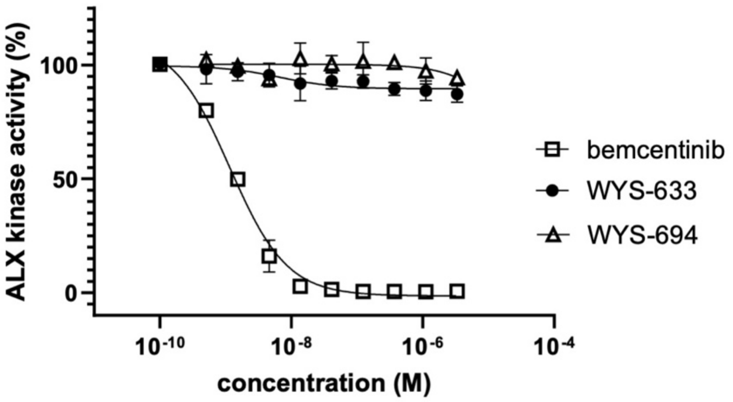


**Figure S3. Effects of compounds on AXL kinase activity.** The AXL kinase activity in the absence of compound was set as 100%.

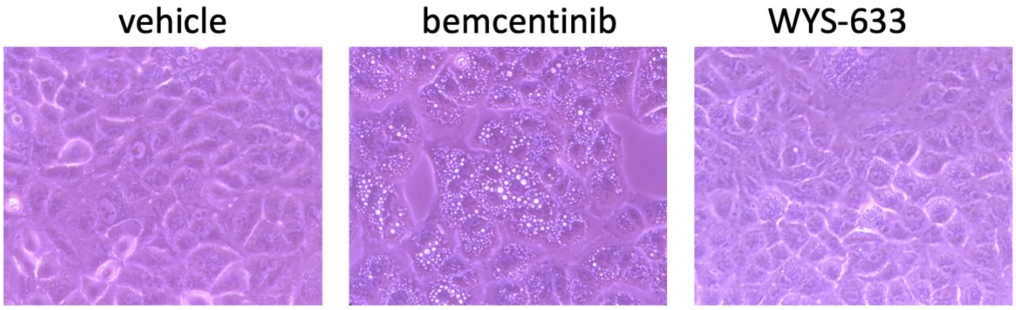


**Figure S4. A549 cell vacuolization.** Cells were incubated with DMSO (vehicle) or 5 μM of compounds for 24h. Phase-contrast images were captured with Echo Revolve. The figure shows representative pictures of at least three independent experiments.

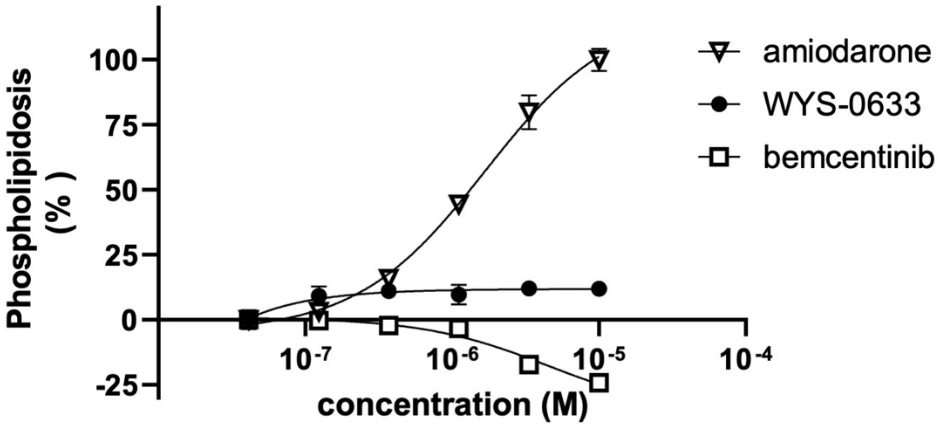


**Figure S5. A549 cell phospholipidosis.** Cells were incubated with DMSO (vehicle) or compounds for 24h. Fluorescence intensities of NBD-PE and Hoechst33342 were measured at wavelengthts of 485/538 or 355/460 (Ex/Em), respectively. Normalized values were calculated by dividing of the NBD-PE value by the Hoechst33342 value and normalized to 10 μM of amiodarone (100%). The graph shows representative concentration curves of three independent experiments. Each data point is the mean ± SD (standard deviation) from three replicates.

| | Bemcentinib | | WYS-633 | |
|---|---|---|---|---|
| | $EC_{50}$ (μM) | n | $EC_{50}$ (μM) | n |
| **SARS-CoV-2** | 0.07 ± 0.04 | 3 | 0.61 ± 0.61 | 7 |
| **BA.2 SARS-CoV-2pp** | 0.16 ± 0.03 | 3 | 0.6 ± 0.3 | 3 |
| **AXL kinase** | 0.00089 ± 0.00001 | 3 | N/A | 3 |

**Table S1. $EC_{50}$ values for bemcentinib and WYS-633 in:** 1) SARS-CoV-2=GFP infected A549-hACE2 cells; 2) BA.2 omicron SARS-CoV-2pp infected HEK-293-hACE2 cells; and 3) AXL kinase enzyme assay. Values are means ± SD of data NA= No activity

| | Bemcentinib | | WYS-633 | | WYS-694 | |
|---|---|---|---|---|---|---|
| | $IC_{50}$ (µM) | n | $IC_{50}$ (µM) | n | $IC_{50}$ (µM) | n |
| **A549-hACE2** | 5.4 ± 1.4 | 4 | 11.1 ± 8.9 | 8 | ND | |
| **HEK-293-hACE2** | 4.2 ± 0.6 | 5 | 8.3 ± 1.1 | 9 | 22.1 ± 12.5 | 6 |
| **HELA-hDPP4** | 2.7 ± 0.3 | 2 | 9.8 ± 0.1 | 2 | ND | |

Values are means ± SD of data from 2 to 9 independent experiments (ND: not determined).

**Table S2. $IC_{50}$ values for Bemcentinib, WYS-633 and WYS-694** on cell viability in ACE2-expressing A549 cells, in ACE2-expressing HEK293 cells or in DPP4-expressing HELA cells.
